# Cognitive functions relate to the decision-dependent component of football skill in youth academy players

**DOI:** 10.64898/2026.11.24.768799

**Authors:** Artur Czeszumski, Mateusz Ludwiczak, Kayla Kolff, Jakub Grządzielewski, Julia Jasińska, Klaudiusz Hirsch, Farbod Nosrat Nezami

## Abstract

Talent identification in football relies increasingly on objective tests. However, these tests capture physical performance better than decision-making, that is, perceiving a game situation and choosing an action. It also remains unclear whether general cognitive functions relate to decision-making on the pitch, partly because studies usually judge skill by playing level or coach ratings, which do not separate decision-making from athletic ability. To address this gap, we tested 79 male academy players aged 12-18 years with a computerized cognitive battery and a football simulator that scores football-specific cognition (decision-making), accuracy, ball control and athletic performance separately. The computerized battery yielded six domain scores: processing speed, executive control, attentional stability, dynamic attention, working memory and cognitive resilience. Processing speed and cognitive resilience correlated with the simulator’s cognition score, whereas executive control and working memory did not. An overall cognitive score correlated with accuracy, the decision-dependent component, but not with athletic performance. Exploratory analyses located this association mainly in passing accuracy. The six cognitive domains explained an additional 22% of the variance in accuracy beyond age, and athletic performance. Therefore, cognitive functions were associated with the decision-dependent component of objectively measured skill, not with its athletic component, which supports adding targeted cognitive tests to academy assessment batteries.

## Introduction

In football, identifying talent is shifting from experience-based judgment toward an objective and multidimensional assessment (Gong et al., 2026). Objective assessments give consistent, comparable information that complements expert judgment and can reduce individual bias in selection (Höner et al., 2021). Such information matters most between the Under-16 (U16) and Under-21 (U21) age groups, when selection decisions have long-term consequences and potential is hardest to judge in players whose current performance is average (Gilson et al., 2025; Tomas et al., 2026). However, objective assessment covers some determinants of performance much better than others.

Objective testing is most advanced for physical performance. Anthropometric, sprint, power and endurance tests follow standardized, validated protocols and dominate academy test batteries (King et al., 2024; Silvino et al., 2024). Technical and tactical skill has no comparable standard. Studies differ in the skills they test (passing, shooting, dribbling, decision-making), in the test setting (isolated drills, video tasks, small-sided games) and in scoring, so results cannot be compared across studies or referenced to age groups (Höner et al., 2021; Leyhr et al., 2018; Machado et al., 2024). Therefore, football clubs still rely on coach ratings and scouting reports, which are open to halo effects – a cognitive bias where one positive trait in a person shapes how others judge that person’s other qualities (Abate Daga et al., 2024). Thus, the part of skill that depends on perceiving a situation and choosing an action is still judged by eye.

Cognitive functions are natural candidates for measuring this perceptual and decision-making part of skill objectively. Coaches and scouts already name cognitive attributes among their selection criteria (Bergkamp et al., 2022; Fuhre et al., 2022), and higher-level players outperform lower-level players on tests of executive functions, the higher-order processes that control behavior, such as inhibition, working memory and cognitive flexibility (Huijgen et al., 2015; Vestberg et al., 2012, 2017). Yet meta-analyses show that this advantage is small on general cognitive tests without sport content and larger on sport-specific tasks, such as video-based decision-making tests (Kalén et al., 2021; Scharfen & Memmert, 2019). Longitudinal data from academy players further show that executive functions develop as in the general population, which questions their value for talent identification (Beavan et al., 2020). This raises the question of whether general cognitive testing belongs in talent identification.

Two features of the evidence keep this question open. First, the criterion is usually coarse: playing level, coach ratings or match statistics, none of which separates the decision-dependent from the athletic components of skill. Second, general and football-specific cognition are rarely measured in the same players (Reinhard et al., 2025), so it is unclear which general functions a football-specific score reflects. Football simulators, such as the SKILLS LAB ARENA (hereafter Arena), now offer a criterion that addresses both points. The Arena delivers balls through a machine, shows game situations on surrounding screens and tracks every action automatically. It returns separate scores for football-specific cognition, accuracy, ball control and athletic performance, with moderate-to-good test–retest reliability in 13-14-year-old elite players (Dolata et al., 2025). As a result, lab-measured cognition can be tested against specific components of objectively measured football skill.

We tackled three main questions. First, which cognitive functions relate to the Arena’s football-specific cognition score (RQ1)? Second, which components of football skill relate to which cognitive functions (RQ2)? Third, does cognition explain variance in skill beyond athletic performance, that is, does it add information that physical testing does not provide (RQ3)? To address these questions, we combined a computerized cognitive battery with the Arena assessment in 79 male academy players aged 12-18 years. Four tasks (choice reaction time, flanker, visual working memory and multiple-object tracking) yielded six domain scores, which we for this study defined as: processing speed, executive control, attentional stability, dynamic attention, working memory and cognitive resilience. Cognitive resilience was characterized as preserved performance under visual distraction; how each score was derived is described in the Methods. All analyses accounted for age, because players were 12-18 years old. For the first question, we expected positive associations for processing speed, dynamic attention and cognitive resilience. We based these expectations on the manufacturer’s description of the football-specific cognition score as visual perception and speed of action, and on the moving teammates and opponents shown in the exercises. For the second question, we contrasted accuracy, which in the passing exercises depends on perceiving the situation and selecting a target, with athletic performance, and tested the difference directly. The answers show whether, and for which part of skill, cognitive testing adds information to an academy’s test battery.

## Methods

### Participants

79 male youth football players from the academy of a Polish first-division professional club took part. Players were 12-18 years old at the Arena assessment (*M* = 14.8 years, *SD* = 1.5; Table 1). All players in the academy’s U14 to U19 teams were invited to take part in the study. Players who were injured or ill at the time were excluded from the study. All 79 players completed both assessments, and there were no missing data on any study variable. Data were pseudonymized before analysis. The study was conducted in accordance with the Declaration of Helsinki. Written informed consent was obtained from parents or legal guardians, and assent from all players according to club policy.

**Table 1.** Participant Characteristics and Scores.

| **Variable** | **All (*n* = 79)** | **ProAssessment (*n* = 49)** | **ProAssessment Talent (*n* = 30)** |
|---|---|---|---|
| Age, years | 14.8 (1.5) | 15.7 (1.1) | 13.3 (0.7) |
| Age range, years | 12-18 | 13-18 | 12-14 |
| *Cognitive domains* |  |  |  |
| Processing speed | 69.1 (10.1) | 71.3 (9.0) | 65.5 (10.7) |
| Executive control (Flanker) | 55.9 (15.2) | 54.9 (15.7) | 57.6 (14.5) |
| Attentional stability | 68.1 (9.4) | 69.9 (8.9) | 65.1 (9.4) |
| Dynamic attention (MOT) | 54.9 (13.7) | 54.8 (13.4) | 55.1 (14.3) |
| Working memory | 51.9 (18.8) | 55.3 (17.7) | 46.3 (19.4) |
| Cognitive resilience | 46.9 (13.9) | 49.5 (12.3) | 42.7 (15.6) |
| *Arena components* |  |  |  |
| Arena cognition | 45.6 (13.5) | 45.4 (10.4) | 46.0 (17.5) |
| Accuracy | 53.3 (9.7) | 51.0 (8.9) | 57.1 (10.0) |
| Ball control | 56.3 (11.5) | 51.2 (9.0) | 64.5 (10.3) |
| Athletics | 65.1 (13.5) | 61.0 (10.7) | 71.7 (15.0) |

Values are M (SD) unless stated otherwise. Cognitive and Arena scores are in points on 1–100 scales (higher = better). Arena scores are shown as raw points; because the two protocols are not on a common scale, all analyses used Arena scores z-standardized within protocol (see Table S1).

### Study Design and Setting

This cross-sectional study combined a computer-based cognitive assessment with an objective assessment of football skill in the Arena simulator (https://skills-lab.com/). Cognitive assessments took place between June and August 2026 and Arena assessments between July and September 2026. The dates of testing were determined largely by the teams’ and each player’s schedules: to avoid interfering with other activities and responsibilities, the researchers set the dates in consultation with the coaches. The study is reported following the STROBE guidelines for cross-sectional studies (von Elm et al., 2007).

### AI Use Declaration

During the preparation of this work, the authors used Claude (Anthropic, Claude Opus 5.5) for language editing and grammar refinement, assistance with writing and debugging data-analysis code, and minor formatting/polishing of figures. Following the use of this tool, the authors thoroughly reviewed, verified, and edited all text, code, and visual elements, and take full responsibility for the accuracy, integrity, and content of the published article.

### Measures

#### Cognitive assessment

##### Apparatus and general procedure

Cognitive functions were assessed with a custom computerized test battery written in Python 3.10 with PsychoPy 2025.2.4 (Peirce et al., 2019). Testing took place in a quiet room in which up to 12 players were assessed simultaneously at separate workstations. The battery ran in full-screen mode on identical computers connected to 21-inch monitors. Players adjusted their workstations to their preference and sat approximately 80 cm from the screen. Stimuli were shown on a light-gray background. The software scaled all stimuli in proportion to the height of the display (26.2 cm); sizes and distances are reported below in degrees of visual angle for the 80 cm viewing distance. Responses were given with the arrow keys of the keyboard in the first three tasks, using only the index finger of the dominant hand, and with the mouse in the fourth task.

All instructions were standardized and presented on screen in Polish or English, together with illustrated examples of each task. They asked players to respond as quickly and accurately as possible and placed particular emphasis on speed in the two tasks scored on reaction time (RT). The battery comprised four tasks administered in a fixed order (Figure 1): a four-choice reaction-time task (processing speed), an arrow version of the Eriksen flanker task (inhibitory control and selective attention; (Eriksen & Eriksen, 1974)), a visual working-memory task (color change detection; (Luck & Vogel, 1997)), and a multiple-object-tracking task (MOT; dynamic visuospatial attention; (Pylyshyn & Storm, 1988)). Each task began with self-paced instructions, which players confirmed they had understood, followed by a short practice block without distractors (five trials; ten trials in the working-memory task, where instructions and practice were repeated automatically if fewer than five responses were correct). Players could repeat the instructions and practice on request. The main block then started after a 10-s countdown. Mandatory 60s rest breaks separated the tasks, and a session lasted approximately 45 min. In all tasks, a white central fixation cross was shown during the intertrial interval (600-1,000 ms, uniformly distributed). No accuracy feedback was given; when no response was registered within the response window of the first three tasks, a “too slow” message appeared for 1 s. Stimulus condition, response, accuracy, and RT were logged for every trial.

**Figure 1.**
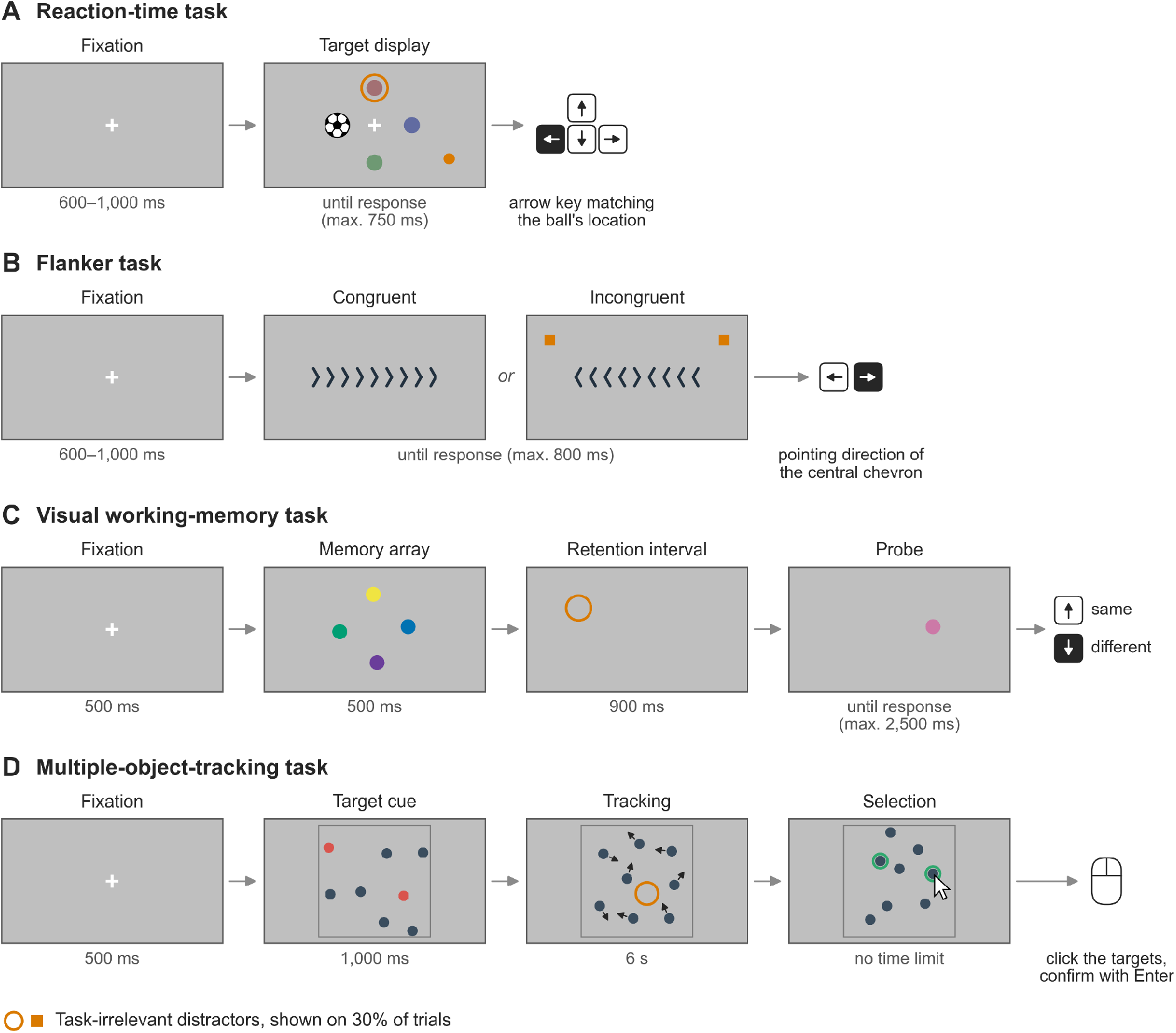
Tasks of the Cognitive Battery. Gray rectangles represent the screen; labels above them name the phases of a trial and labels below them give their durations. (A) Reaction-time task: a football appeared at one of four locations together with three colored discs, and players pressed the arrow key matching the football’s location. (B) Flanker task: players reported the pointing direction of the central chevron with the left or right arrow key while ignoring the flanking chevrons, which pointed in the same direction (congruent) or the opposite direction (incongruent). (C) Visual working-memory task: players memorized an array of two to five colored discs and, after a blank retention interval, judged whether a single probe disc had the same color as the item previously shown at that location (up arrow) or a different color (down arrow). (D) MOT: two to six target discs were highlighted in red among 8-18 identical discs, all discs then moved for 6s (arrows indicate motion and were not shown), and players clicked on the discs they had tracked (selected discs were outlined in green). In C and D, an intertrial interval of 600-1,000 ms followed the response (not shown). Dark keys mark the correct response for the example shown. Orange shapes illustrate the task-irrelevant distractors that appeared on a random 30% of trials in every task: flickering shapes during the stimulus display in A and B, a brief flash during the retention interval in C, and repeated flashes during tracking in D. Stimuli are not drawn to scale.

##### Reaction-time task

Four stimuli appeared simultaneously around the central fixation cross: one to its left, one to its right, one above it, and one below it, each centered 5.6° from fixation. One of them was a picture of a football (3.7° wide), which served as the target; the other three were colored discs (muted blue, red, and green; 2.4° in diameter). Players pressed the arrow key corresponding to the location of the football. The display remained on screen until a response was made or for a maximum of 750 ms. The task comprised 120 trials, with the target appearing 30 times at each location in random order. A reminder to respond as fast as possible was displayed for 1.8 s every 15-20 trials.

##### Flanker task

A horizontal row of nine chevrons was presented until response or for a maximum of 800 ms. Each chevron was 0.5° wide and 1.6° high, and the centers of neighboring chevrons were 1.4° apart, so the whole row was 11.7° wide. Players indicated the pointing direction of the central chevron with the left or right arrow key and ignored the four flanking chevrons on each side, which pointed in the same direction (congruent) or the opposite direction (incongruent). The row appeared at one of five vertical positions selected at random on each trial (at the screen center, or 2.8° or 4.7° above or below it), so that the vertical position of the target was unpredictable. The task comprised 240 trials in random order (120 congruent and 120 incongruent; 120 left-pointing and 120 right-pointing targets), with a 60s rest break after 120 trials and the same periodic speed reminder as in the reaction-time task.

##### Visual working-memory task

Each trial began with a fixation cross (500 ms), followed by a memory array of two, three, four, or five colored discs (1.3° in diameter) shown for 500 ms, a blank retention interval of 900 ms, and a single probe disc presented at the location of one randomly selected memory item until response or for a maximum of 2,500 ms. The memory items were equally spaced on an imaginary circle centered on fixation (radius 4.7°), and each item was shifted at random by up to 0.7° horizontally and vertically. Their colors were drawn without replacement from a fixed palette of eight colors. With a probability of 0.5, the probe had the same color as the item previously shown at that location; otherwise it had a different color drawn at random from the remaining seven, which could be a color shown elsewhere in the array. Players pressed the up arrow key for “same” and the down arrow key for “different”. This mapping was displayed at the bottom of the screen during the probe and repeated in a 3-s reminder every 10-15 trials. Key presses within the first 180 ms after probe onset were ignored. The task comprised 120 trials, 30 at each set size, blocked by set size in ascending order, with 60-s rest breaks between blocks.

##### MOT

After a fixation cross (500 ms), 8-18 identical dark discs (0.9° in diameter) appeared at random, non-overlapping positions inside a square frame (16.7° × 16.7°), and 2-6 of them were highlighted in red for 1,000ms to mark them as targets. All discs then became identical and moved for 6s along straight paths in random directions, each at a constant speed drawn from a uniform distribution between 3.4°/s and 5.6°/s. Discs bounced off the borders of the frame, moved independently of one another, and could overlap. Eye movements were not restricted. When the motion stopped, the mouse cursor appeared at the screen center and players clicked on the discs they believed to be the targets. Clicking a selected disc again cancelled the selection. Players had to select exactly as many discs as there were targets and then confirmed their response with the Enter key; there was no time limit. The task comprised 40 trials at five levels of tracking load presented in ascending order: 2 targets among 8 discs (4 trials), 3 among 10 (6 trials), 4 among 12 (14 trials), 5 among 14 (12 trials), and 6 among 18 (4 trials).

##### Distractor manipulation

In every task, 30% of the main-block trials contained task-irrelevant visual distractors (36 of 120 trials in the reaction-time task, 72 of 240 in the flanker task, 36 of 120 in the working-memory task, and 12 of 40 in the MOT task). These trials were selected at random, independently of the other trial factors; the remaining trials are referred to as baseline trials. Distractors were orange shapes with abrupt onsets at task-irrelevant locations. In the reaction-time task, two shapes flickered at 4 Hz for as long as the stimuli were displayed: a ring (radius 1.8°) around one of the three non-target discs, and a small disc (radius 0.8°) at a random screen location that changed with every flicker cycle. In the flanker task, two squares (side length 1.1°) flickered at 5 Hz in one of two arrangements: in the upper left and upper right of the screen (9.3° to either side of the screen center and 5.6° above it), or directly above and below the chevron row, 6.3° from it. In the second arrangement, the square farther from the screen center was partly or fully off screen whenever the row was not at the screen center. In the working-memory task, a ring (radius 1.5°) was flashed for 150 ms at a random location in the middle of the retention interval (405-555 ms after offset of the memory array). In the MOT task, a ring (radius 1.7°) was flashed for 120 ms at a random location inside the frame every 1.2s during tracking (five flashes per trial). Distractors were not mentioned in the instructions.

##### Domain scores

Performance was summarized in six domain scores ranging from 1 to 100, with higher scores indicating better performance. Each raw performance index x was rescaled linearly between two fixed anchors, which were defined a priori to represent poor and excellent performance: score = 100 × (x − poor anchor) / (excellent anchor − poor anchor). The result was truncated to the range 1-100. The scores are therefore criterion-referenced; they are not standardized against the present sample or against age norms. The first five scores were computed from baseline trials only. RT indices were based on correct responses, and no RT trimming was applied. Processing speed was derived from the reaction-time task and combined the median correct RT, accuracy, and the omission rate (proportion of trials without a response). Executive control reflected the flanker interference effect, that is, the difference between incongruent and congruent trials in median correct RT and in accuracy. Attentional stability reflected intraindividual variability of correct RTs pooled across the reaction-time and flanker tasks. It combined the coefficient of variation (SD/mean) with a lapse rate, defined as the proportion of correct responses slower than the player’s median RT plus 2.5 scaled median absolute deviations, plus the proportion of trials without a response. Dynamic attention was the effective number of tracked targets in the MOT task, K = T × (H − FA), where T is the number of targets, H the proportion of targets selected, and FA the proportion of non-targets selected; K was computed for each trial, bounded between 0 and T, and averaged across trials. Working memory was the capacity estimate K = S × (H − FA) (Cowan, 2001; Rouder et al., 2011), where S is the set size, H the proportion of “different” responses on change trials, and FA the proportion of “different” responses on no-change trials; K was computed for each set size, bounded between 0 and S, and averaged across set sizes, weighted by the number of trials. Cognitive resilience quantified how well performance was preserved on distractor trials relative to baseline trials. For each task, distractor costs were computed (relative slowing of the median correct RT and loss of accuracy in the reaction-time and flanker tasks; relative reduction of K and loss of accuracy in the working-memory task; relative reduction of K in the MOT task), converted to scores on which the absence of a cost corresponds to 100, and averaged with equal weights across the four tasks. These six domain scores were the cognitive variables of this study.

#### Football skills

Football skill was assessed in the Skills Lab Arena, a football simulator in which automated ball machines deliver balls to the player and laser-based tracking, high-speed cameras and software record every action under identical conditions for all players. Players completed one of two assessment protocols: ProAssessment (*n* = 49; 13-18 years) or ProAssessment Talent (hereafter Talent), a protocol for younger players with easier settings in three exercises (*n* = 30; 12-14 years; see below). Players were assigned to a protocol by team, younger teams to ProAssessment Talent, and older to ProAssessment.

Both protocols are standardized sequences of exercises in which the player receives balls from the ball machines, interprets football situations shown on the screens surrounding the pitch, identifies a target according to a predefined rule and responds with a dribble, pass or shot that is tracked automatically. Each protocol comprises 12 exercises, two of which are performed twice, giving 13 runs with 140 delivered balls plus a timed dribbling course (Table S9). The first exercise, a one-two passing exercise with 10 slow, low balls, is designated by the manufacturer as a warm-up that familiarizes the player with the ball deliveries, the displays and the passing targets. It is followed by two dribbling exercises that end with a pass to a moving teammate shown on the screens (10 balls each); a half-turn passing exercise, in which the player scans over the shoulder before receiving the ball, turns through 180° and passes to a teammate (two runs of 20 balls); an overview exercise, in which the player identifies the screen on which teammates have a numerical advantage and passes to one of them (two runs of 20 balls); four shooting-accuracy exercises, in which the player takes one touch and shoots at a target in a lower or an upper corner of the goal with the right or the left foot (6 balls each); two shooting-power exercises, in which the player shoots 3 stationary balls with each foot from 11 m for maximum ball speed; and a sprint course, in which the player dribbles around four cones as fast as possible.

The two protocols differ in three exercises. In the half-turn passing exercise of ProAssessment, teammates and opponents move in both runs, and the second run has more opponents and faster and lifted balls delivered by four instead of two ball machines; in Talent, teammates and opponents are static in the first run, and in the second run the teammates move, the opponents remain static and the ball delivery is unchanged. The shooting-accuracy exercises require a minimum shot speed of 70 km/h in ProAssessment and 50 km/h in Talent. The sprint course comprises two laps in ProAssessment and one lap in Talent. All other exercises and settings are identical in the two protocols. According to the manufacturer, the assessment is not designed to assess tactical behavior, endurance, long-distance sprinting or duels.

The software reports eight basic parameters on a 1-100 point scale (higher = better): passing accuracy, shooting accuracy, shooting speed, agility, dribbling, first touch, two-footedness and cognition, the last described by the manufacturer as visual perception and speed of action. The basic parameters are grouped into category scores. Because the manufacturer does not publish these weights, we regressed each category score on the basic parameters in our sample (n = 79); the category scores were reproduced almost exactly by fixed combinations of the basic parameters (*R*² = 0.988-0.999): accuracy by equal parts of passing and shooting accuracy; ball control by two-thirds first touch and one-third dribbling; and athletics by agility and shooting speed (weights ≈ 0.68 and 0.36). In 13-14-year-old elite players, Arena tasks showed intersession intraclass correlation coefficients of 0.75-0.89 for time-based and 0.71-0.91 for accuracy-based tasks (Dolata et al., 2025).

The analyses used four Arena scores, hereafter the Arena components: cognition (hereafter Arena cognition), accuracy, ball control and athletics. We regarded cognition and accuracy as the components that depend most on perceiving the situation and selecting an action, and athletics (agility and shooting speed) as the athletic component. For accuracy, this applies mainly to its passing half: in the passing exercises the player must first identify the target teammate on the screens, whereas in the shooting-accuracy exercises the target is fixed and performance depends more on technique. Ball control (first touch and dribbling) combines technical and athletic demands.

#### Data Preparation

To check whether the two Arena protocols are on a common scale, each of the four raw Arena component scores and the cognitive score (defined below) were regressed on protocol and age, with HC3 standard errors. These analyses showed that the protocols are not on a common scale (see Results and Table S1). Arena scores were therefore *z*-standardized within protocol, and protocol was entered as a covariate in all models. The six cognitive domains were only modestly intercorrelated (median Spearman ρ = 0.21, range −0.07 to 0.66; Cronbach’s α of the *z*-scored domains = 0.68), so each domain was analyzed separately. Where a single summary of cognition was required (the specificity test in RQ2 and Figure 3), we used the unweighted mean of the six *z*-scored domains (cognitive score). Results with the overall composite score of the battery software (undisclosed weights for each domain) and with the first principal component of the six domains are reported as sensitivity analyses.

#### Statistical Analysis

Associations were estimated with Spearman partial correlations controlling for age, recorded in completed years at the Arena assessment, and protocol (ProAssessment vs ProAssessment Talent). Spearman correlations were chosen because several scores were skewed or bounded. All variables were rank-transformed, the ranks of each variable of interest were residualized on the ranks of the covariates, and the residuals were correlated. p values were obtained from the t distribution with *n* − 4 degrees of freedom. Ninety-five percent confidence intervals (CIs) were bias-corrected and accelerated (BCa) bootstrap intervals (Efron, 1987) based on 5,000 resamples, with players drawn with replacement within each protocol.

To describe development and to justify the age adjustment, each cognitive domain was also regressed on age with heteroscedasticity-consistent (HC3) standard errors, with Holm correction across the six domains. The change in cognitive score per year was expressed in standard-deviation units, and mean domain scores by age group are reported descriptively. A two-tailed α of 0.05 was used throughout.

Analyses were run in Python 3.10 with NumPy (Harris et al., 2020), pandas (McKinney, 2010), SciPy (Virtanen et al., 2020), statsmodels (Seabold & Perktold, 2010) and Pingouin (Vallat, 2018), and figures were drawn with Matplotlib (Hunter, 2007). The analyses for each research question are described below.

### RQ1: Which cognitive functions relate to football-specific cognition?

We correlated each of the six domains with Arena cognition and controlled the familywise error rate across the six tests with the Holm procedure (Holm, 1979). On theoretical grounds we expected positive associations for processing speed, dynamic attention and cognitive resilience.

### RQ2: Which components of football skill relate to which cognitive functions?

We correlated the six domains with accuracy, ball control and athletics and controlled the false discovery rate across the 18 tests with the Benjamini–Hochberg procedure (Benjamini & Hochberg, 1995). A significant correlation with one component and a non-significant correlation with another does not show that the two correlations differ, so specificity was tested directly. We estimated the difference between the cognitive score’s partial correlations with accuracy and with athletics, Δρ = ρ(accuracy) − ρ(athletics), with a BCa bootstrap CI in which both correlations were computed in the same resamples; a CI excluding zero indicated a reliable difference. The same difference was computed for each domain as a descriptive breakdown without correction.

To evaluate the absence of an association between the cognitive score and athletics, we used two one-sided tests (TOST) of equivalence (Lakens, 2017) with bounds of ρ = ±0.30, approximately the smallest correlation detectable with 80% power in this sample. The tests used Fisher’s *z* with the standard error of Bonett and Wright (2000) for Spearman correlations and were repeated for each domain as part of the descriptive breakdown. For the cognitive score we also computed a default Bayes factor for the corresponding Pearson partial correlation (Ly et al., 2016), with a stretched beta prior of width κ = 1, and the sample size reduced by the number of covariates, and interpreted BF_10_ < 1/3 as moderate evidence for the absence of an association (Jeffreys, 1961).

### RQ3: Does cognition explain skill beyond athletic performance?

We fitted a hierarchical linear regression with accuracy as the outcome, age, protocol and athletics in step 1, and the six cognitive domains as one block in step 2; continuous variables were standardized. The contribution of cognition was tested with the *F*-test of the change in *R*² (Δ*R*²), with Cohen’s *f*² as the effect size (Cohen, 1988), and adjusted *R*² is reported for both steps. Because heteroscedasticity can bias conventional standard errors, the block was also tested with a Wald test based on HC3 standard errors (Long & Ervin, 2000; MacKinnon & White, 1985), and HC3-based 95% CIs are reported for the standardized coefficients (β). Because the domains overlap, individual coefficients describe unique contributions and are less precise than the block test, which was the test of RQ3. Multicollinearity was assessed with variance inflation factors (VIF), normality of residuals with the Shapiro–Wilk test (Shapiro & Wilk, 1965), heteroscedasticity with the Breusch–Pagan test (Breusch & Pagan, 1979), and influential cases with Cook’s distance (> 4/*n*; (Cook, 1977).

### Robustness and Sensitivity Analyses

The three research question conclusions were re-examined in three alternative analyses: (a) Pearson instead of Spearman partial correlations; (b) excluding players with an extreme value (|robust *z*| > 3.5, based on the median and the median absolute deviation; (Leys et al., 2013)) on any analyzed score, with Arena scores evaluated within protocol; and (c) the ProAssessment protocol only, with age as the only covariate. The same multiplicity corrections were applied within each alternative, and CIs were based on Fisher’s *z*. A conclusion was considered robust when its significance decision and sign were unchanged. Results for the Talent protocol alone (*n* = 30) are reported descriptively. Whether the association between the cognitive score and accuracy differed between the protocols was tested with Fisher’s *z* test for independent correlations, using Bonett–Wright variances. The specificity and incremental-validity tests were repeated with the battery’s composite score and with the first principal component of the six *z*-scored domains in place of the cognitive score (for RQ3, as a single predictor in step 2). In an exploratory analysis, the six domains were correlated with all eight basic Arena parameters, with Benjamini–Hochberg correction across the 48 tests.

## Results

All 79 players had complete data (Table 1). Players tested with the ProAssessment Talent protocol were younger than those tested with ProAssessment (*M* = 13.3 vs 15.7 years). At equal age, Talent players scored higher than ProAssessment players on accuracy (by 10.6 points, 95% CI [4.4, 16.9], *p* < 0.001), ball control (16.4 points [8.5, 24.3], *p* < 0.001) and athletics (16.9 points [6.6, 27.2], *p* = 0.001), but not on Arena cognition (−0.4 points [−10.4, 9.6], *p* = 0.937) or on the cognitive score (0.00 *SD* [−0.40, 0.40], *p* = 0.996; Table S1). Because younger players would not be expected to outperform older players on the same test, this pattern indicates that the protocols are not on a common scale and supports within-protocol standardization.

The cognitive score increased by 0.13 *SD* per year of age (95% CI [0.03, 0.24], *p* = 0.011; Table S2). After Holm correction, processing speed (*b* = 2.65 points per year, 95% CI [1.04, 4.26], *p*_Holm_ = 0.008) and cognitive resilience (*b* = 3.15 [1.02, 5.28], *p*_Holm_ = 0.019) increased with age. The increase in working memory did not survive correction (*b* = 3.20 [0.67, 5.74], *p*_Holm_ = 0.053), and executive control, attentional stability and dynamic attention showed no reliable age trend (means by age group in Table S3). Age was therefore controlled in all analyses.

### Cognitive Functions and Football-Specific Cognition

After adjustment for age and protocol, Arena cognition correlated with cognitive resilience (ρ = 0.37, 95% CI [0.15, 0.55], *p*_Holm_ = 0.006) and processing speed (ρ = 0.34 [0.14, 0.52], *p*_Holm_ = 0.013; Figure 2, first column; Table S4). The correlations with attentional stability (ρ = 0.28 [0.05, 0.49], *p*_Holm_ = 0.054) and dynamic attention (ρ = 0.26 [0.03, 0.45], *p*_Holm_ = 0.074) were positive but did not survive correction. Executive control (ρ = 0.05 [−0.17, 0.28]) and working memory (ρ = −0.06 [−0.27, 0.17]) were unrelated to Arena cognition (both *p*_Holm_ = 1.000). Thus, two of the three expected associations (processing speed and cognitive resilience) were confirmed.

**Figure 2.**
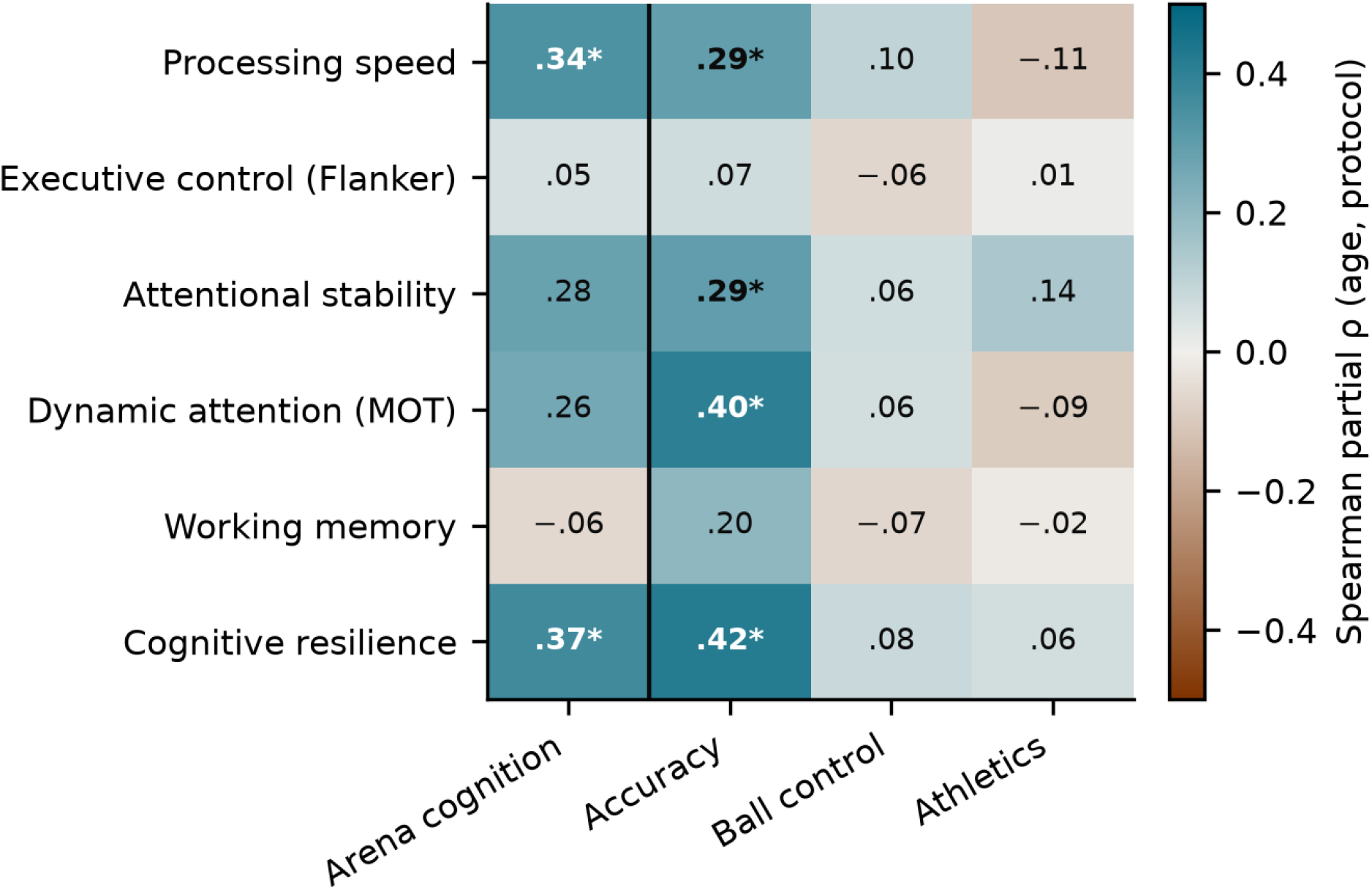
Partial Correlations Between the Cognitive Domains and the Arena Components. Spearman partial correlations controlling for age and protocol (*n* = 79); Arena scores were standardized within protocol. Bold values marked with an asterisk were significant after correction: Holm correction across six tests for Arena cognition (RQ1) and Benjamini–Hochberg correction across 18 tests for accuracy, ball control and athletics (RQ2). Blue indicates positive and red negative correlations. 95% CIs and *p* values are given in Table S4. The vertical line separates the RQ1 column from the RQ2 columns.

### Skill Components and Cognitive Functions

Of the 18 correlations between the six domains and accuracy, ball control and athletics, four survived Benjamini–Hochberg correction, all with accuracy (Figure 2; Table S4): cognitive resilience (ρ = 0.42, 95% CI [0.21, 0.60], *p*_BH_ = 0.002), dynamic attention (ρ = 0.40 [0.20, 0.57], *p*_BH_ = 0.002), attentional stability (ρ = 0.29 [0.09, 0.48], *p*_BH_ = 0.048) and processing speed (ρ = 0.29 [0.02, 0.51], *p*_BH_ = 0.048). Working memory (ρ = 0.20 [−0.01, 0.41], *p*_BH_ = 0.292) and executive control (ρ = 0.07 [−0.15, 0.29], *p*_BH_ = 0.672) were not related to accuracy. No domain correlated with ball control or athletics (all |ρ| ≤ 0.14, all *p*_BH_ ≥ 0.635).

The cognitive score correlated with accuracy (ρ = 0.45, 95% CI [0.26, 0.62]) but not with athletics (ρ = −0.01 [−0.24, 0.20]), and the difference between the two correlations was reliable (Δρ = 0.46, 95% CI [0.16, 0.77]; Figure 3). The correlation with athletics was statistically equivalent to zero within ±0.30 (TOST *p* = 0.005), and the Bayes factor provided moderate evidence for the absence of an association (BF_10_ = 0.15). For single domains, the accuracy–athletics difference was reliable for dynamic attention (Δρ = 0.49 [0.20, 0.77]), processing speed (Δρ = 0.40 [0.07, 0.69]) and cognitive resilience (Δρ = 0.36 [0.03, 0.67]), but not for working memory (Δρ = 0.22 [−0.08, 0.50]), attentional stability (Δρ = 0.15 [−0.15, 0.45]) or executive control (Δρ = 0.06 [−0.26, 0.39]; Table S5). The correlation with athletics was equivalent to zero for five of the six domains; the exception was attentional stability (TOST *p* = 0.080).

**Figure 3.**
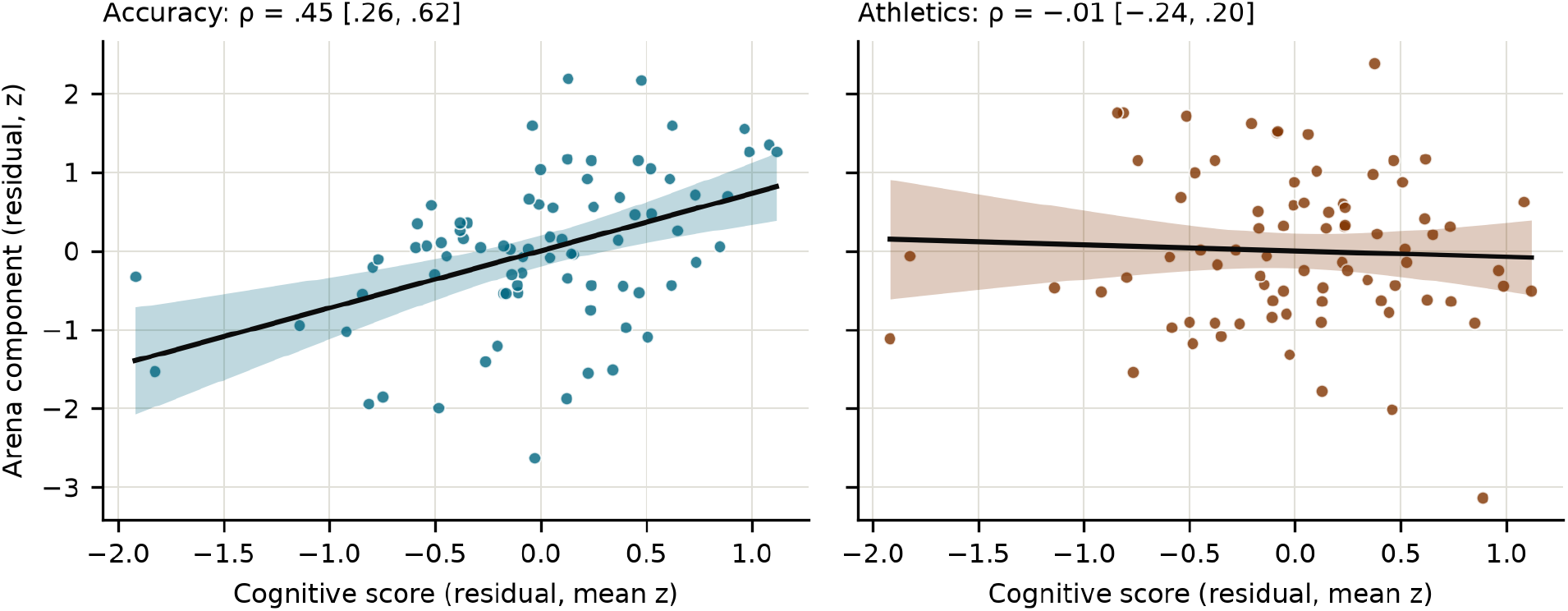
Cognitive Score Against Arena Accuracy and Athletics After Adjustment for Age and Protocol. Partial residuals after regressing each variable on age and protocol (*n* = 79). The cognitive score is the unweighted mean of the six *z*-scored cognitive domains; Arena scores are *z*-scores within protocol. Lines show linear fits with 95% confidence bands. Panel titles give Spearman partial correlations with 95% BCa CIs. The difference between the two correlations is Δρ = 0.46, 95% CI [0.16, 0.77].

### Cognition Beyond Athletic Performance

Age, protocol and athletics explained 5% of the variance in accuracy (*R*² = 0.05, adjusted *R*² = 0.01; Table 2). Adding the six cognitive domains increased the explained variance to 27% (*R*² = 0.27, adjusted *R*² = 0.18), Δ*R*² = 0.22, *F*(6, 69) = 3.52, *p* = 0.004, *f*² = 0.31. The HC3-robust test of the block gave the same conclusion, *F*(6, 69) = 4.00, *p* = 0.002. Athletics did not predict accuracy (β = −0.04, 95% CI [−0.24, 0.17]). Among the domains, only dynamic attention made a unique contribution whose CI excluded zero (β = 0.22 [0.02, 0.43], *p* = 0.030; Figure 4). Cognitive resilience had the largest coefficient (β = 0.29 [−0.13, 0.70], *p* = 0.174), but it overlapped with the other domains (VIF = 2.93). All VIFs were below 3, residuals were consistent with normality (Shapiro–Wilk *p* = 0.311) and homoscedasticity (Breusch–Pagan *p* = 0.359), and six players had a Cook’s distance above 4/*n* (Figure S1).

**Table 2.** Hierarchical regression of Arena accuracy on age, protocol, athletics and the six cognitive domains.

| **Predictor** | **β** | **95% CI** | ***p*** | **VIF** |
|---|---|---|---|---|
| Age | 0.19 | [−0.24, 0.62] | 0.379 | 2.72 |
| Protocol (Talent vs ProAssessment) | 0.50 | [−0.17, 1.16] | 0.143 | 2.46 |
| Athletics | −0.04 | [−0.24, 0.17] | 0.732 | 1.11 |
| Processing speed | −0.02 | [−0.41, 0.36] | 0.901 | 1.93 |
| Executive control (Flanker) | −0.03 | [−0.26, 0.21] | 0.832 | 1.15 |
| Attentional stability | 0.08 | [−0.19, 0.35] | 0.567 | 2.05 |
| Dynamic attention (MOT) | 0.22 | [0.02, 0.43] | 0.030 | 1.34 |
| Working memory | 0.08 | [−0.17, 0.32] | 0.538 | 1.30 |
| Cognitive resilience | 0.29 | [−0.13, 0.70] | 0.174 | 2.93 |

Coefficients of the step 2 model (*n* = 79). Continuous variables were standardized; accuracy and athletics are *z*-scores within protocol; protocol was coded 0 = ProAssessment, 1 = ProAssessment Talent. CIs and *p* values are based on HC3 standard errors. Step 1 (age, protocol, athletics): *R*² = 0.05, adjusted *R*² = 0.01. Step 2 (adding the six domains): *R*² = 0.27, adjusted *R*² = 0.18; Δ*R*² = 0.22, *F*(6, 69) = 3.52, *p* = 0.004; HC3 Wald test of the block *F*(6, 69) = 4.00, *p* = 0.002. VIF = variance inflation factor.

**Figure 4.**
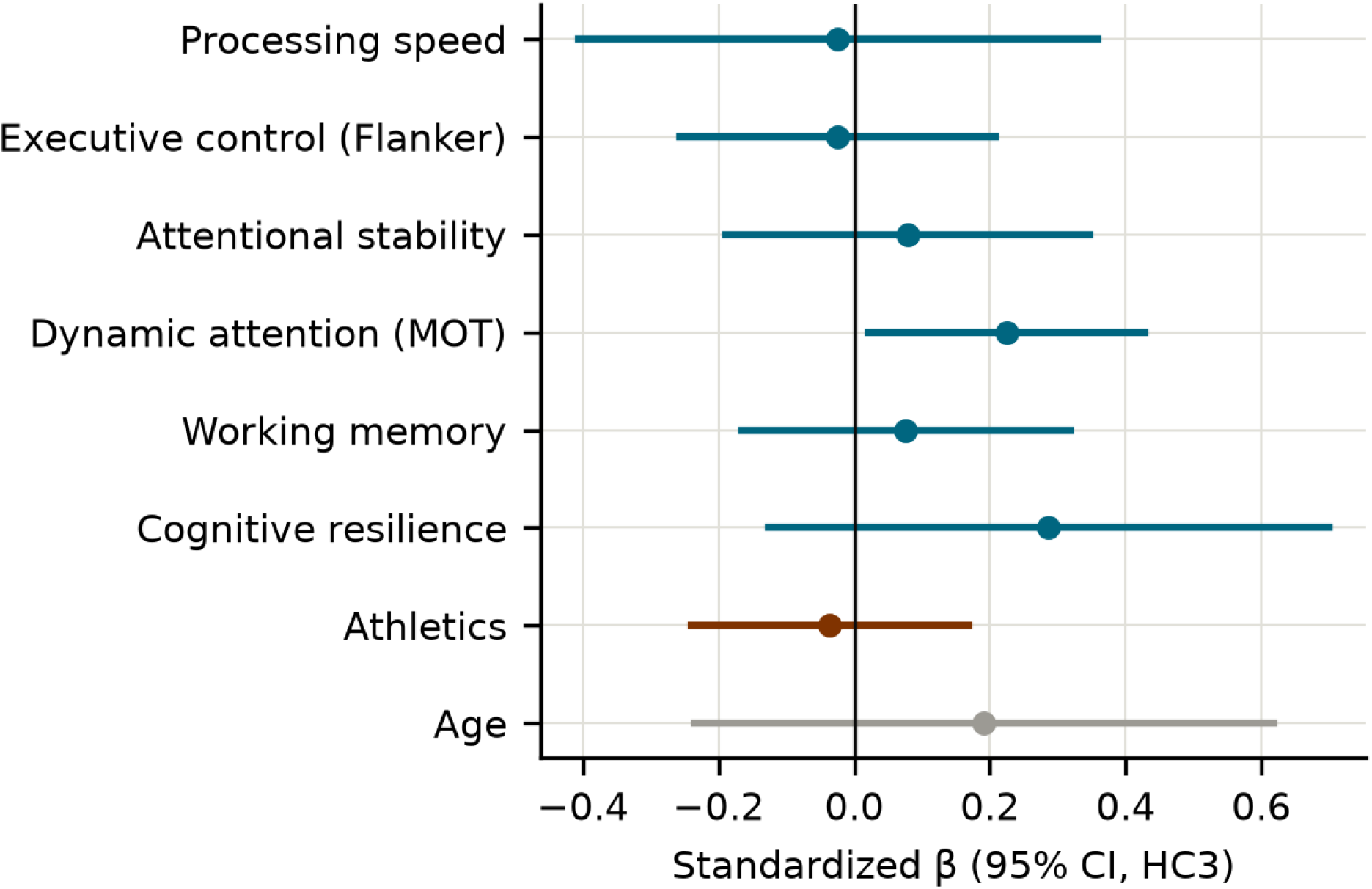
Standardized Coefficients of the Step 2 Model Predicting Arena Accuracy. Points show standardized coefficients (β) with HC3 95% CIs from the model including age, protocol, athletics and the six cognitive domains (*n* = 79). Blue = cognitive domains, orange = athletics, gray = age; the protocol coefficient is reported in Table 2. Together, the six domains added Δ*R*² = 0.22 (*p* = 0.004).

### Robustness and Sensitivity Analyses

With Pearson correlations, and after excluding the five players with an extreme value (*n* = 74), the associations of processing speed and cognitive resilience with Arena cognition (RQ1), the associations of dynamic attention and cognitive resilience with accuracy, the specificity difference and the RQ3 increment were all unchanged (Table S6). The associations of processing speed (*r* = 0.25) and attentional stability (*r* = 0.27) with accuracy were not significant with Pearson correlations, and the association of attentional stability was also not significant after excluding extreme values (ρ = 0.26); these two associations are therefore less robust. Conversely, attentional stability (*r* = 0.32) and dynamic attention (*r* = 0.29) correlated significantly with Arena cognition when Pearson correlations were used.

In the ProAssessment protocol alone (*n* = 49), the RQ1 associations were replicated (processing speed ρ = 0.46, cognitive resilience ρ = 0.39; attentional stability, ρ = 0.40, was also significant). The associations with accuracy, however, were smaller (ρ = 0.27–0.31 for the four domains that were significant in the full sample) and not significant after correction, and neither the specificity difference (Δρ = 0.24) nor the RQ3 increment (Δ*R*² = 0.21) was significant. The correlation between the cognitive score and accuracy was 0.33 among ProAssessment players and 0.63 among Talent players; this difference was not significant (*z* = 1.53, *p* = 0.125). With 49 players, the ProAssessment-only analysis had 80% power only for |*r*| ≥ 0.39. Among Talent players (descriptive), dynamic attention (ρ = 0.51) and cognitive resilience (ρ = 0.58) correlated with accuracy more strongly than in the full sample.

The conclusions did not depend on how the six domains were summarized (Table S7). With the battery’s composite score, the correlations were ρ = 0.39 with accuracy and ρ = −0.06 with athletics (Δρ = 0.45 [0.13, 0.75]), and the composite explained Δ*R*² = 0.16 beyond age, protocol and athletics (*p* < 0.001). With the first principal component (43% of variance; weights 0.05 for executive control and 0.32–0.56 for the other domains), the correlations were ρ = 0.49 and 0.01 (Δρ = 0.48 [0.18, 0.78]; Δ*R*² = 0.19, *p* < 0.001). The corresponding values for the cognitive score were Δρ = 0.46 and, as a single predictor, Δ*R*² = 0.18 (*p* < 0.001).

In the exploratory analysis of the eight basic Arena parameters, four of the 48 correlations survived correction (Table S8): passing accuracy correlated with dynamic attention (ρ = 0.43, 95% CI [0.24, 0.61], *p*_BH_ = 0.003) and cognitive resilience (ρ = 0.42 [0.23, 0.59], *p*_BH_ = 0.003), and Arena cognition with cognitive resilience (ρ = 0.37, *p*_BH_ = 0.017) and processing speed (ρ = 0.34, *p*_BH_ = 0.031). Correlations with shooting accuracy were weaker (all ρ ≤ 0.22) and none survived correction, so the accuracy associations were carried mainly by passing accuracy.

## Discussion

This study examined whether and to what extent lab-measured cognitive functions relate to objectively measured football skills in academy players. We found that processing speed (the speed and accuracy of choice responses in the reaction-time task) and cognitive resilience (preserved performance under visual distraction) correlated with the Arena’s football-specific cognition score (described by the manufacturer as visual perception, speed of action and decision-making), whereas executive control (flanker interference) and working memory (change-detection capacity) did not. In addition, cognition was related to accuracy but not to ball control or athletics. The cognitive score correlated more strongly with accuracy than with athletics. The six cognitive domains explained an additional 22% of the variance in accuracy beyond age, protocol and athletics. Therefore, in these players, cognition was tied to the decision-dependent component of skill, and not to its athletic component.

The lab battery and the Arena’s cognition score converged on speed and on resistance to distraction. The manufacturer describes this score as visual perception and speed of action, so its correlation with processing speed was expected. The correlation with cognitive resilience suggests that the score also reflects how well a player keeps performing when irrelevant events compete for attention. Attentional stability and dynamic attention pointed in the same direction but did not survive correction, so our expectation for dynamic attention was not confirmed. Executive control and working memory, for which we had not predicted associations, were unrelated to Arena cognition. We interpret the null result for executive control with caution. Flanker interference is a difference score, and such scores often rank individuals unreliably (Hedge et al., 2018). The confirmed correlations were moderate, so the lab-based and Arena measures of cognition overlap but are not interchangeable. This is consistent with the view that general tests capture only part of sport-specific cognition (Kalén et al., 2021). Our findings therefore show that general processing speed and resistance to distraction, but not interference control or working-memory capacity, track the football-specific cognition score of this simulator.

Of the three skill components, cognition related only to accuracy. Accuracy correlated with cognitive resilience, dynamic attention, attentional stability and processing speed, although the last two associations were less robust in sensitivity analyses. None of the six cognitive domains correlated with ball control or athletics. For athletics, equivalence and Bayesian tests supported the absence of an association with the cognitive score. We tested this specificity directly, because a significant correlation with accuracy and a non-significant correlation with athletics do not by themselves show that the two correlations differ (Gelman & Stern, 2006; Nieuwenhuis et al., 2011). Exploratory analyses located the effect in passing accuracy (dynamic attention ρ = 0.43; cognitive resilience ρ = 0.42), not in shooting accuracy (all ρ ≤ 0.22). This pattern can be explained by the task demands: in the Arena’s passing exercises, the player must identify the target teammate among the players shown on the screens, whereas the shooting targets are fixed. Some studies that use playing level as the criterion find little or no link with general cognition (Furley et al., 2026; Reinhard et al., 2025). Playing level reflects all components of skill at once, so an association with one component, such as passing accuracy, can be diluted by components unrelated to cognition, such as athletic ability; a criterion measured at the level of components can reveal such associations. In our data, cognition was therefore specific to the component of skill that requires selecting a target, not to ball handling or athletic output.

The cognitive domains also explained accuracy beyond athletic performance. Age, protocol and athletics together explained 5% of the variance in accuracy; adding the six cognitive domains raised this to 27%. Athletics itself did not predict accuracy. Among the domains, only dynamic attention made a unique contribution whose confidence interval excluded zero; cognitive resilience had the largest coefficient, but its interval included zero because it overlapped with the other domains. A limitation is that athletics in the Arena meant agility with the ball and shooting speed, not sprint, jump or endurance performance. Within the simulator, cognition therefore explained variance in decision-dependent skill that athletic performance did not.

Cognitive scores were higher in older players. The cognitive score increased per year, driven by processing speed and cognitive resilience. Working memory showed a similar slope, but this trend did not survive correction (Holm-corrected p = 0.053) and should be interpreted with caution. Executive control, attentional stability and dynamic attention showed no reliable age trend. Faster processing in older adolescents matches normative development (Kail, 1991; Luna et al., 2004) and longitudinal data from academy players (Beavan et al., 2020). The flat profile for dynamic attention differs from the increase between 13 and 15 years reported for three-dimensional tracking in 404 players (Zhang et al., 2025) and may reflect our smaller sample or our two-dimensional task. Because these are cross-sectional differences between age groups, they describe the sample and not individual development.

The findings were not equally strong in the two protocols. The associations with Arena cognition were replicated when the ProAssessment protocol (n = 49) was analyzed alone. In this protocol, however, the associations with accuracy were smaller and, like the specificity difference (Δρ = 0.24) and the increment (ΔR² = 0.21), not significant. Among Talent players the cognitive score correlated with accuracy at ρ = 0.63, against 0.33 among ProAssessment players. This difference was itself not significant (p = 0.125), and the ProAssessment subsample had 80% power only for |r| ≥ 0.39. Protocol and age are also confounded: Talent players were 12–14 years old and completed an easier protocol, whereas ProAssessment players were mostly older and completed a harder one. A weaker link in ProAssessment could reflect older age, the harder protocol or chance, and our data cannot separate these explanations. Therefore the pooled estimates need replication, especially in older players.

In academy football players, lab-measured cognitive functions were associated with the decision-dependent component of objectively measured skill, mainly passing accuracy, and not with its athletic component. They explained variance in accuracy that age and athletic performance did not. In sum, assessment batteries that aim to capture this component of skill should include cognitive measures matched to the decision-making demands of the game.

## Practical Implications

For practice, the results indicate both what cognitive testing can add to an academy’s battery and where its limits lie. A 45-minute computerized battery captured variance in accuracy, mainly passing accuracy, that athletic testing missed, which supports cognitive tests as a complement to physical and technical assessment. The functions worth targeting are those that are related to accuracy in this study, namely dynamic attention, cognitive resilience and processing speed, rather than executive functions as a block. The battery is more accessible than a simulator but cannot replace it, because the two capture only partly overlapping aspects of performance. Two steps should precede any use in selection. Longitudinal studies must show that these scores predict skill development, selection or the transition to senior football, which earlier longitudinal work questions for executive functions (Beavan et al., 2020). Intervention studies must show whether training these functions improves passing; so far, three-dimensional tracking training improved passing decisions in one small trial (Romeas et al., 2016) but not in a larger follow-up (Romeas et al., 2025). Until then, we recommend that cognitive scores are better suited to describing players than to selecting them.

## Limitations

Four further limitations qualify these conclusions. First, the design is cross-sectional, so the associations do not show that cognition predicts later skill, selection or progression. Second, the criterion was a simulator, not match performance or coach ratings. Its scores are proprietary composites, and its passing exercises present targets on screens, so some shared variance may reflect screen-based visual processing common to both assessments. Third, the cognitive battery was custom-built: the reliability of its six scores was not estimated, and cognitive resilience is derived from the same tasks as the other five. Fourth, the sample was 79 male players from one academy, which gives 80% power only for correlations of about 0.30 or larger, and we recorded chronological age although biological maturity varies widely at a given age (Malina et al., 2015). These limits define the claim: cognition and decision-dependent skill were associated in this sample, under these measures.

## Author contributions

A.C.: Conceptualization, Data curation, Formal analysis, Investigation, Methodology, Supervision, Validation, Visualization, Writing – original draft, Writing – review & editing. M.L.: Writing – original draft, Writing – review & editing. K.K.: Writing – review & editing. J.G.: Investigation. J.J.: Investigation, K.H.: Investigation. F.N.N.: Writing – review & editing.

